# Label-free prediction of three-dimensional fluorescence images from transmitted light microscopy

**DOI:** 10.1101/289504

**Authors:** Chawin Ounkomol, Sharmishtaa Seshamani, Mary M. Maleckar, Forrest Collman, Gregory R. Johnson

## Abstract

Understanding cells as integrated systems is a challenge central to modern biology. While different microscopy approaches may be used to probe diverse aspects of biological organization, each method presents limitations which ultimately restrict a view into unified cellular organization. For example, while fluorescence microscopy can resolve subcellular structure in living cells, it is expensive, slow, and can damage cells. Here, we present a label-free method for predicting 3D fluorescence directly from transmitted light images and demonstrate that it can be used to generate multi-structure, integrated images. We then demonstrate that this same method can be used to predict immunofluorescence from electron micrograph inputs, extending the method to a wider range of bioimaging applications.

---

The various imaging methods currently used to capture details of cellular organization all present restrictions with respect to expense, spatio-temporal resolution, and sample perturbation. Fluorescence microscopy permits imaging of specific proteins and structures of interest by labeling them specifically; but it requires advanced instrumentation and time-consuming sample preparation. Critically, samples are subject to significant phototoxicity and photobleaching, creating a tradeoff between data quality and time scales available for live cell imaging^1,2^. Furthermore, the number of simultaneous fluorescent tags is restricted by both spectrum saturation and cell health, limiting the number of parallel labels that can be imaged together. Transmitted light microscopy, e.g., bright-field, phase, DIC, etc., in contrast, is relatively low-cost and label/dye-free, with greatly reduced phototoxicity^3^ and simplified sample preparation. Although valuable information about cellular organization is apparent in transmitted light images; they lack the clear contrast of fluorescence labeling, a limitation also present in other widely used microscopy modalities. Electron micrographs, for example, contain a rich set of biological detail about subcellular structure, but often require tedious expert interpretation. A method that could combine the clarity of fluorescence microscopy with the relative simplicity and modest cost of other imaging techniques would present a groundbreaking tool for biological insight into the integrated organization of subcellular structures.

Convolutional neural networks (CNNs) capture non-linear relationships over large spatial areas of images, resulting in vastly improved performance for image recognition tasks as compared to classical machine learning methods. Here, we present a CNN-based tool, employing a U-Net architecture^4^ (Supplementary Fig. 1, see Methods) to model the relationships between distinct but correlated imaging modalities, and show the efficacy of this tool for predicting corresponding fluorescence images directly from both 3D transmitted light live cell images and 2D electron micrographs, alone.

The label-free imaging tool learns the relationship between 3D transmitted light (bright-field and DIC) and fluorescence live cell images corresponding to several major subcellular structures (, e.g., cell membrane, nuclear envelope, nucleoli, DNA, and mitochondria; Fig. 1a). The resultant model can then both predict a 3D fluorescence image from a completely new transmitted light input, and both combine predictions for a variety of subcellular structures can be combined, effectively enabling multi-channel, integrated fluorescence imaging from a single transmitted light input (Fig. 1d, e). Utility is not limited to transmitted light microscopy; the method can similarly be used to predict 2D immunofluorescence (IF) images directly from never-before-seen electron micrographs (EM) to clearly highlight distinct subcellular structures and the registration of conjugate conjugate multi-channel fluorescence data with EM^5^ (Fig. 2).

**Figure 1:**
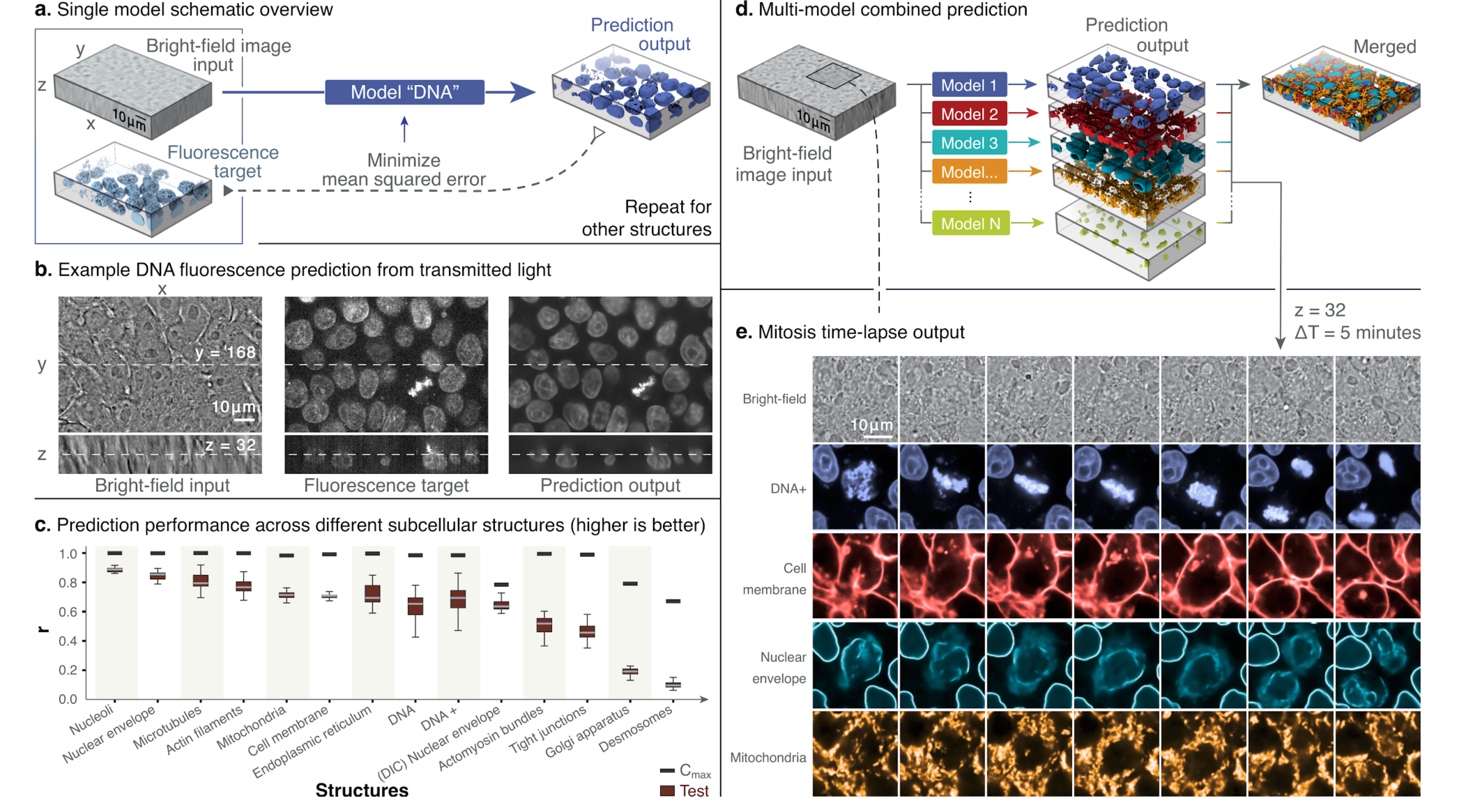
Label-free imaging tool pipeline and application using 3D transmitted light-to-fluorescence models. a) Given transmitted light and fluorescence image pairs as input, the model is trained to minimize the mean squared error (MSE) between the fluorescence ground truth and output of the model. b) Left to right, an example of a 3D input transmitted light image, a ground-truth confocal DNA fluorescence image, and a tool prediction. c) Distributions of the image-wise correlation coefficient (*r*) between ground truth (target) and predicted test images derived from the indicated subcellular structure models. Each target/predicted image pair in the test set is a point in the resultant *r* distribution; the 25th, 50th and 75th percentile image pairs are spanned by the box for each indicated structure, with whiskers indicating the last data points within the 1.5x interquartile range of the lower and upper quartiles. For complete description of the structure labels, see Methods. Black bars indicate maximum correlation between the target image and a theoretical, noise-free image (Cmax; for details see Methods). d) Individual subcellular structure models are applied to the same input and combined to predict multiple structures. e) Localization of DNA (blue), cell membrane (red), nuclear envelope (cyan) and mitochondria (orange) as predicted for time lapse transmitted light (bright-field) input images taken at 5-minute intervals (center z-slice shown); a mitotic event with stereotypical reorganization of subcellular structures is clearly evident. All results shown here are obtained from new transmitted light images not used during model training.

**Figure 2:**
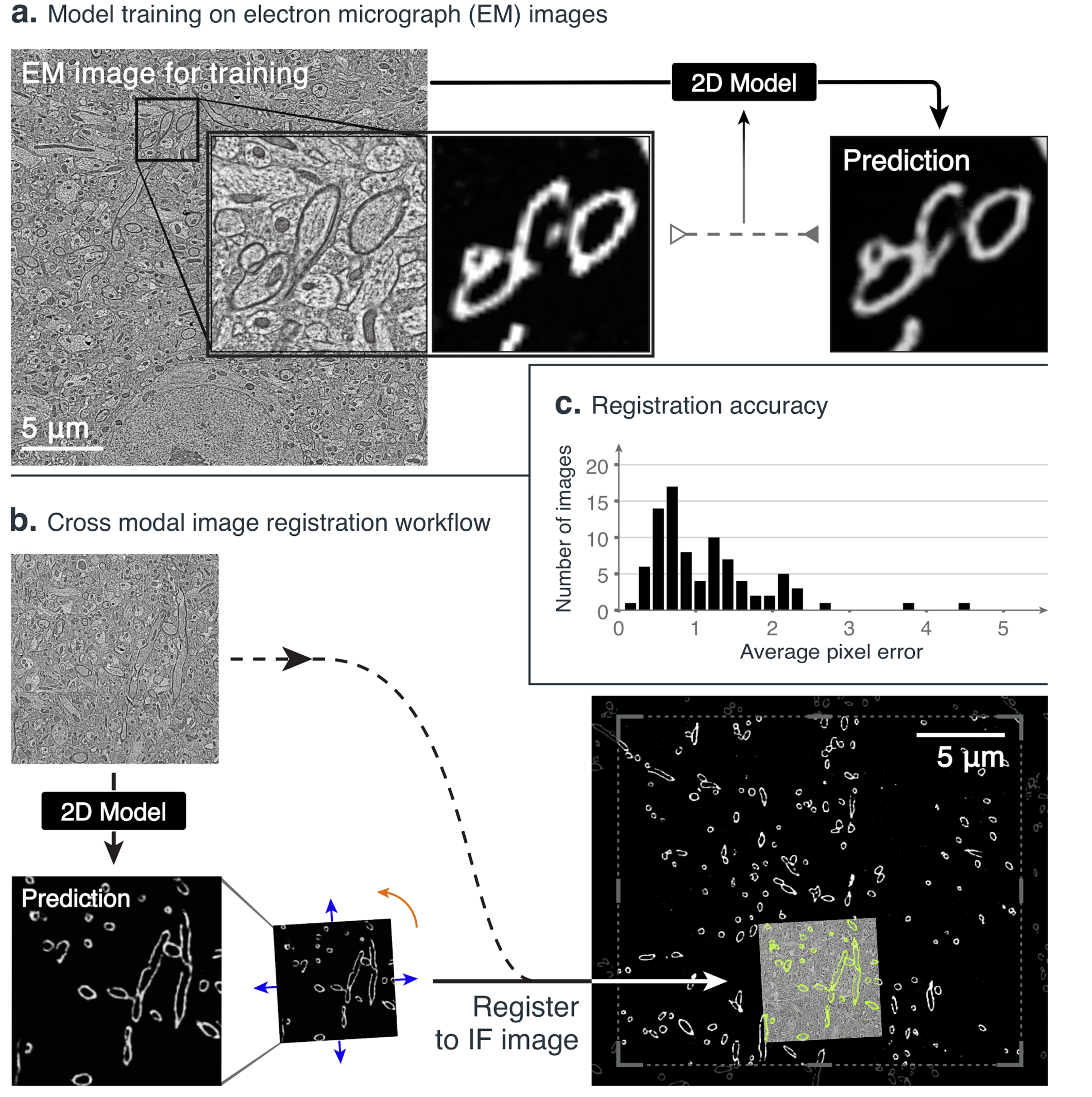
Label-free imaging tool facilitates 2D automated registration across imaging modalities. We first train a model to predict a 2D myelin basic protein immunofluorescence image (MBP-IF) from a 2D electron micrograph (EM) and then register this prediction to automate cross-modal registration. a) An example EM image with a highlighted subregion (left), the MBP-IF image corresponding to the same subregion (middle), and the label-free imaging tool prediction of the same subregion given only the EM image as input (right). b) The EM image of the subregion to be registered (top left) is passed through the trained 2D model to obtain a prediction for the subregion (bottom left), which is then registered to MBP-IF images within a larger field of view (bottom right) (see Methods for details). Only a 20 *μ*m × 20 *μ*m region from the 204.8 *μ*m × 204.8 *μ*m MBP-IF search image is shown; predicted and registered MBP-IF are overlaid (in green) together with the EM image. c) Histogram of average distance between automated registration and manual registration as measured across 90 test images, in units of pixels of MBP-IF data. This distribution has an average of 1.16 ± 0.79 px, where manual registrations between two independent annotators differed by 0.35 ± 0.2 px.

This method can train a model to learn a relationship between two imaging modalities for the structure of interest given only spatially registered pairs of images, from a relatively small image set (30 image pairs per structure for 3D transmitted light-to-fluorescence, and 40 for 2D EM-to-fluorescence; see Methods). The biological detail that can be observed in predictions varies among subcellular structures modeled. However, in the case of the 3D transmitted light-to-fluorescence models, for example, predicted images appear structurally similar to ground truth fluorescence images in 3D. As examples, nuclear structures are well-resolved: images produced by the DNA model (Fig. 1b) depict well-formed and separated nuclear regions as well as finer detail, including chromatin condensation just before and during mitosis, and the nuclear envelope model predictions (Supplementary Fig. 2) provide high-resolution localization and 3D morphology. The nucleoli model also resolves the precise location and morphology of individual nucleoli (Supplementary Fig. 2). Models for several other structures also perform favorably upon visual inspection as compared to ground-truth images; the mitochondria model correctly identifies the regions of cells with high numbers of mitochondria as well as regions which are more sparsely populated. In general, individual mitochondria visible in the fluorescence data are also observable in predictions (Supplementary Fig. 2). While predictions for microtubules and the endoplasmic reticulum do not resolve individual filaments or detailed morphology, respectively, these models successfully capture broader 3D localization of the structures.

The transmitted light-to-fluorescence models’ performance was quantified via the Pearson’s correlation coefficient on 20 predicted images and their corresponding ground truth fluorescence images image pairs (Fig. 1c) from independent test sets for each model (see Methods). A theoretical upper bound was determined for ideal model performance based upon an estimate of the signal-to-noise ratio (SNR) of the fluorescence images used for training (see Methods). The models’ performance for each structure is well-bounded by this limit (Fig. 1c).

Given these promising results, we trained the model predicting DNA with an extended training procedure (DNA+) to evaluate whether the outcome could be improved with additional training images and iterations. As expected, the performance improved as measured by an increase in Pearson’s correlation, and the images showed qualitatively, better clarity of the sub-nuclear structure and precision of the predictions around mitotic cells (Fig. 1c, Supplementary Fig. 2, Methods) Most critically, all of these details can be observed together in a 3D integrated multi-channel prediction derived from a single transmitted light image (Fig. 1e, Supplementary Fig. 2 and Supplementary Video). Examples for all fourteen subcellular labeled structure models’ predictions on a test set (new images that model did not see during training) can be found in Supplementary Fig. 2.

Transforming one imaging modality to another proves useful for a variety of imaging challenges; just as the label-free tool can be used to predict 3D integrated fluorescence images from a single transmitted light input, the 2D IF images predicted from EM (Fig. 2) can be used to facilitate automatic registration of conjugate multi-channel fluorescence data with EM. Array tomography data^5^ of ultrathin brain sections uses EM and ten channels of IF images (including the structure myelin basic protein, MBP-IF) obtained from the same sample, but from two different microscopes, and thus EM and IF images are not natively spatially aligned. While EM and corresponding images of other modalities can be registered by hand through identification of corresponding locations and fitting a similarity transformation, resulting in multi-channel conjugate EM images^5,6,7^, manual registration is tedious and time-consuming. We trained a 2D version of the label-free tool on manually registered pairs of EM and MBP-IF images and then evaluated the tool in the scenario of registering an EM image (15 *μ*m × 15 *μ*m) to a target MBP-IF image covering a much larger area (204.8 *μ*m × 204.8 *μ*m) (Fig. 2a). For this, EM images (of regions not used in the training set) were input to the model to predict corresponding MBP-IF images (Fig. 2a) and conventional intensity-based matching techniques (see Methods) were then used to register each MBP-IF prediction (and thus the EM image) to the target MBP-IF image (Fig. 2b), which converged successfully on 86 of 90 image pairs. The average distance between automated and manual registrations for 90 test image pairs was 1.16 ± 0.79 px, whereas manual registrations between two independent annotators differed by 0.35 ± 0.2 px. To the authors’ knowledge, this is the first successful attempt to automate this registration process via conventional statistical image registration techniques, suggesting that the label-free tool’s utility can be extended to diverse imaging modalities and a variety of additional downstream image processing tasks.

We next determined that the models for individual structures, trained solely on static images, can be used to predict temporal fluorescence image sequences. To do this, we applied several subcellular structure transmitted light-to-fluorescence models to a single transmitted light 3D time-series (covering 95 minutes at 5-minute intervals; Fig. 1e, Supplementary Video 1). In addition to simultaneous visualization of several subcellular structures, characteristic dynamics of mitotic events, including the reorganization of the nuclear envelope and cell membrane, are evident in the predicted multi-channel time-series (Fig. 1e). Time series acquired with a similar 5-minute acquisition interval, under imaging conditions used to acquire training data, but with three-channel spinning disk fluorescence, reveal both obvious bleaching artifacts and changes in cellular morphology and health after 10-15 minutes (data not shown). This phototoxicity, which often occurs in extended, multi-label live cell time-series fluorescence imaging on the hiPSCs used for this study, shows the challenges in obtaining this information from normal fluorescence time lapse imaging. While many strategies exist to minimize this photodamage (i.e. oxygen scavenging, reduced laser power and exposure, advanced microscopy techniques^2^, machine learning driven denoising^8^), all present compromises with respect to ease, image quality, information content, and fidelity. This method avoids these trade-offs and directly produces time-series predictions for which no fluorescence imaging ground truth is available, greatly increasing the timescales over which some cellular processes can be visualized and measured.

Of course, our method has inherent limitations and is not well suited for all applications. Because models must learn a relationship between distinct but correlated imaging modes, predictive performance is contingent upon the existence of this association. In the case of desmosomes or actomyosin bundles, for example, model performance for the presented training protocol was comparatively poor, presumably due to a weaker association between transmitted light and fluorescence images of these structures (Fig. 1c, Supplementary Fig. 2). The quality and quantity of training data will also influence accuracy of the model predictions, although this relationship is highly nonlinear in tested cases (for DNA model performance, we see diminishing returns between 30 and 60 images; Supplementary Fig. 3). Performance between models additionally varies with 2D and 3D information: predictions using a 2D DNA model to predict z-slices selected from 3D images show artifacts between predicted z-slices and a decrease in correlation between ground truth and predicted images (Supplementary Fig. 5; see Methods), suggesting that the 3D interference pattern is valuable for predicting subcellular organization.

We cannot assess *a priori* how models will perform in biological contexts for which there are very few or no examples in training or testing data. Specifically, models pre-trained using one cell type (i.e. hiPSC) do not perform as well when applied to inputs with drastically different cellular morphologies (Supplementary Fig. 4). We compared the predictions from the DNA+ model (trained on hiPSC images) to those of a model trained on images of DNA-labeled HEK-293 kidney-phenotype cells^9^ when applied to both hiPSC and HEK-293 test images. While gross image features are comparable, in this example, prediction performance of morphological details improves markedly when the model is trained on the same cell type. When a pre-trained model is used to predict a fluorescent DNA label in very different cell types, like cardiomyocytes or HT-1080 fibroblast-phenotype cells^10^ (Supplementary Fig 4, see Methods) we see a similar reduction in predictive performance.

Furthermore, predictions from inputs acquired with imaging parameters identical to those used to compose a model training set will provide the most accurate results when compared to ground truth data; while we successfully trained fluorescence models using both bright-field and DIC modalities, using bright-field images as input to a model trained on DIC images resulted in poorer model performance. More subtle parameters can also influence performance: we observed a decrease in model accuracy when predicting fluorescence images from input transmitted light stacks acquired with a shorter inter-slice interval (~0.13 s) than that in training data (~2.2 s) (data not shown). Ultimately, when evaluating the utility of predicted images, the context for which those images will be used must be considered. For instance, DNA or nuclear membrane predictions may potentially have sufficient accuracy for application to downstream nuclear segmentation algorithms, but microtubule predictions would not be effective for assaying rates of microtubule polymerization (Fig. 1e, Supplementary Fig. 2). Finally, it is important to note that there may not be a direct quantitative link between the predicted intensity of a tagged structure and the protein levels.

The label-free methodology presented here has wide potential for use in many biological imaging fields. Primarily, it is possible to reduce or even eliminate routine capture of some images in existing imaging and analysis pipelines, permitting the same throughput in a far more efficient and cost-effective manner. Notably, data used for training requires no manual annotation, little to no pre-processing, and relatively small numbers of paired examples, drastically reducing the barrier to entry associated with some machine learning approaches. Areas where this approach may prove of particular value include image-based screens where cellular phenotypes can be detected via expressed fluorescent labels^11^, pathology workflows requiring specialized labels that identify specific tissues^12^, long time series observation of single cells^1^, tissues, or organism-level populations where more expensive instrumentation is not available^2^. Recent related work convincingly demonstrates that 2D whole-cell antibody stains can be predicted from transmitted light^13^. Although distinct from the present study in method and scope, this work supports the conclusion that similar techniques can be applied to a wide variety of biological studies, as is the case for automatic cross-modal registration of conjugate multi-channel fluorescence data with EM shown here. The presented tool is additionally promising in cases wherein generating a complete set of simultaneous ground-truth labels is not feasible, as is the case for the live cell time-series imaging example presented. Finally, our tool permits the generation of integrated images by which multi-dimensional interactions among cellular components can be investigated. This implies exciting potential for probing coordination of subcellular organization as cells grow, divide, and differentiate, and signifies a new opportunity for understanding structural phenotypes in the context of disease modeling and regenerative medicine. More broadly, the presented work suggests an opportunity for a key new direction in biological imaging research: the exploitation of imaging modalities’ indirect but learnable relationships to visualize biological features of interest with ease, low cost, and high fidelity.

## Methods

### 3D live cell imaging

The 3D light microscopy data used to train and test the presented models consists of z-stacks of colonies of human embryonic kidney cells (HEK-293)^9^, human fibrosarcoma cells (HT-1080)^10^, genome-edited human induced pluripotent stem cell (hiPSC) lines^10^ expressing a protein endogenously tagged with either mEGFP or mTagRFP that localizes to a particular subcellular structure^14^, and hiPSC-derived cardiomyocytes differentiated from the former. The EGFP-tagged proteins and their corresponding structures are: alpha-tubulin (microtubules), beta-actin (actin filaments), desmoplakin (desmosomes), lamin B1 (nuclear envelope), fibrillarin (nucleoli), myosin IIB (actomyosin bundles), sec61B (endoplasmic reticulum), STGAL1 (Golgi apparatus), Tom20 (mitochondria) and ZO1 (tight junctions). The cell membrane was labelled by expressing RFP tagged with a CAAX motif.

#### Tissue Culture

hiPSCs, HEK-293 cells, or HT-1080 cells were seeded onto Matrigel-coated 96-well plates at densities specified below. The cells were stained on the days they were to be imaged, first by incubation in their imaging media with 1x NucBlue (Hoechst 33342, ThermoFisher) for 20 min. hiPSCs were then incubated in imaging media with 1x NucBlue and 3x CellMask (ThermoFisher) for 10 min, whereas HEK-293 and HT-1080 cells were then incubated in imaging media with 1x NucBlue and 0.5x CellMask for 5 min. The cells were washed with fresh imaging media before imaging.

For hiPSCs, the culture media was mTeSR1 (Stem Cell Technologies) with 1% Pen-Strep. The imaging media was phenol-red-free mTeSR1 with 1% Pen-Strep. Cells were seeded at a density of ~2500 cells per well and were imaged 4 days after initial plating. For HEK-293 cells, the culture media was DMEM with GlutaMAX (ThermoFisher), 4.5 g/L D-Glucose, 10% FBS, and 1% antibiotic-antimycotic. The imaging media was phenol-red-free DMEM/F-12 with 10% FBS and 1% antibiotic-antimycotic. Cells were seeded at a density of 13 to 40 thousand cells per well and were imaged 1 to 2 days after initial plating. For HT-1080 cells, the culture media was DMEM with GlutaMAX, 15% FBS, and 1% Pen-Strep. The imaging media was phenol-red-free DMEM/F-12 with 10% FBS and 1% Pen-Strep. Cells were seeded at a density of 2.5 to 40 thousand cells per well and were imaged 4 days after initial plating.

CAAX-tagged hiPSCs were differentiated to cardiomyocyte phenotype by seeding onto Matrigel-coated 6-well tissue culture plates at a density ranging from 0.5–2 × 10^6^ cells per well in mTeSR1 supplemented with 1% Pen-Strep, 10 *μ*M ROCK inhibitor (Stem Cell Technologies), and 1 *μ*M CHIR99021 (Cayman Chemical). The following day (designated day 0), directed cardiac differentiation is initiated by treating the cultures with 100 ng/mL ActivinA (R&D) in RPMI media (Invitrogen) containing 1:60 diluted GFR Matrigel (Corning), and insulin-free B27 supplement (Invitrogen). After 17 hours (day 1), cultures are treated with 10 ng/mL BMP4 (R&D systems) in RPMI media containing 1 *μ*M CHIR99021 and insulin-free B27 supplement. At day 3, cultures are treated with 1 *μ*M XAV 939 (ToCris) in RPMI media supplemented with insulin-free B27 supplement. On day 5, the media is replaced with RPMI media supplemented with insulin-free B27. After differentiation into cardiomyocytes, cells were grown in RPMI media supplemented with B27. Cardiomyocytes were re-plated at day 12 onto glass-bottom plates coated with PEI/laminin and were imaged on day 43 after initiation of differentiation. The imaging media was phenol-red-free RPMI with B27. Prior to imaging, cells were stained by incubating in imaging media with Nuclear Violet (AAT Bioquest) at a 1:7500 dilution and 1x CellMask for 1 min and then washed with fresh imaging media.

#### Imaging

All cell types were imaged for up to 2.5 h on a Zeiss spinning disk microscope with ZEN Blue 2.3 software and with a 1.25-NA, 100x objective (Zeiss C-Apochromat 100x/1.25 W Corr), with up to four, 16-bit data channels per image: transmitted light (either bright-field or DIC), cell membrane labeled with CellMask, DNA labeled with Hoechst, and EGFP-tagged cellular structure. Respectively, acquisition settings for each channel were: white LED, 50 ms exposure; 638 nm laser at 2.4 mW, 200 ms exposure; 405 nm at 0.28 mW, 250 ms exposure; 488 nm laser at 2.3 mW, 200 ms exposure. The exception were CAAX-RFP-based cell membrane images, which were acquired with a 1.2-NA, 63x objective (Zeiss C-Apochromat 63x/1.2 W Corr), a 561 nm laser at 2.4 mW, and a 200 ms exposure. 100x-objective z-slice images were captured at a YX-resolution of 624 px × 924 px with a pixel scale of 0.108 *μ*m/px, and 63x-objective z-slice images were captured at a YX-resolution of 1248 px × 1848 px with a pixel scale of 0.086 *μ*m/px. All z-stacks were composed of 50 to 75 z slices with an inter-z-slice interval of 0.29 *μ*m. Images of cardiomyocytes contained 1 to 5 cells per image whereas images of other cell types contained 10 to 30 cells per image. Time-series data were acquired using the same imaging protocol as for acquisition of training data but on unlabeled, wild-type hiPSCs at 5 minute intervals for 95 minutes, with all laser powers set to zero to reproduce the inter-z-slice timing of the training images.

### Data for training and evaluation

Supplementary Table 1 outlines the data used to train and evaluate the models based on 3D live cell z-stacks, including train-test data splits. All multi-channel z-stacks were obtained from a database of images produced by the Allen Institute for Cell Science’s microscopy pipeline (see http://www.allencell.org). For each of the 11 hiPSC cell lines, we randomly selected z-stacks from the database and paired the transmitted light channel with the EGFP/RFP channel to train and evaluate models (Fig. 1c) to predict the localization of the tagged subcellular structure. The transmitted light channel modality was bright-field for all but the DIC-to-nuclear envelope model. For the DNA model data, we randomly selected 50 z-stacks from the combined pool all bright-field-based z-stacks and paired the transmitted light channel with the Hoechst channel. The training set for the DNA+ model was further expanded to 540 z-stacks with additional images from the Allen Institute for Cell Science’s database. Note that while a CellMask channel was available for all z-stacks, we did not use this channel because the CAAX-membrane cell line provided higher quality images for training cell membrane models. A single z-stack time series of wild-type hiPSCs was used only for evaluation (Fig. 1e).

For experiments testing the effects of number of training images on model performance (Supplementary Fig. 3), we supplemented each model’s training set with additional z-stacks from the database. Z-stacks of HEK-293 cells were used to train and evaluate DNA models whereas all z-stacks of cardiomyocytes and of HT-1080 cells were used only for evaluation (Supplementary Fig. 4). The 2D DNA model (Supplementary Fig. 5) used the same data as the DNA+ model.

All z-stacks were converted to floating-point and were resized via cubic interpolation such that each voxel corresponded to 0.29 *μ*m × 0.29 *μ*m × 0.29 *μ*m, and resulting images were 244 px × 366 px for 100x-objective images or 304 px × 496 px for 63x-objective images in Y and X respectively and between 50 and 75 pixels in Z. Pixel intensities of all input and target images were z-scored on a per-image basis to normalize any systematic differences in illumination intensity.

### Electron and Immunofluorescence Microscopy

#### Imaging

For conjugate array tomography data^5^, images of 50 ultra-thin sections were taken with a wide-field fluorescence microscope using 3 rounds of staining and imaging to obtain 10-channel immunofluorescence (IF) data (including myelin basic protein, MBP) at 100 nm per pixel. 5 small regions were then imaged with a field emission scanning electron microscope to obtain high resolution electron micrographs at 3 nm per pixel. Image processing steps independently stitched the IF sections and one of the EM regions to create 2D montages in each modality. Each EM montage was then manually registered to the corresponding MBP channel montage with TrakEM2^15^.

#### Data Used for Training and Evaluation

40 pairs of registered EM and MBP montages were resampled to 10 nm per pixel. For each montage pair, a central region of size 3280 px × 3214 px was cut out and used for the resultant final training set. This corresponded to the central region of the montage which contained no unimaged regions across the sections used. Pixel intensities of the images were z-scored. For the registration task, a total of 1500 EM images (without montaging) were used as an input to directly register to the corresponding larger MBP image in which it lies. For this, each EM image was first downsampled to 10 nm per pixel without any transformations to generate a 1500 px × 1500 px image.

### Model architecture description and training

We employed a convolutional neural network (CNN) based on the U-Net architecture^4^ (Supplementary Fig. 1) due to its demonstrated performance in image segmentation and tracking tasks. In general, CNNs are uniquely powerful for image-related tasks (classification, segmentation, image-to-image regression) due to the fact that they are image-translation invariant, learn complex non-linear relationships across multiple spatial areas, circumvent the need to engineer data-specific feature extraction pipelines, and are straightforward to implement and train. CNNs have been shown to outperform other state-of-the-art models in basic image recognition^16^ used in biomedical imaging for a wide range of tasks including image classification, object segmentation^17^, and estimation of image transformations^18^. Our U-Net variant consists of layers that perform one of three convolution types, followed by a batch normalization and ReLU operation. The convolutions are either 3 pixel convolutions with a stride of 1-pixel on zero-padded input (such the input and output of that layer are the same spatial area), 2-pixel convolutions with a stride of 2 pixels (to halve the spatial area of the output), or 2-pixel transposed convolutions with a stride of 2 (to double the spatial area of the output). There are no normalization or ReLU operations on the last layer of the network. The number of output channels per layer are shown in Supplementary Fig. 1. The 2D and 3D models use 2D or 3D convolutions, respectively.

Due to memory constraints associated with GPU computing, we trained the model on batches of either 3D patches (64 px × 64 px × 32 px, YXZ) for light microscopy data or on 2D patches (256 px × 256 px) for conjugate array tomography data, which were randomly subsampled uniformly both across all training images as well as spatially within an image. The training procedure took place in a typical forward-backward fashion, updating model parameters via stochastic gradient descent (backpropagation) to minimize the mean squared error between output and target images. All models presented here were trained using the Adam optimizer^19^ with a learning rate of 0.001 and with beta values of 0.5 and 0.999 for 50,000 mini-batch iterations. We used a batch size of 24 for 3D models and of 32 for 2D models. Running on a Pascal Titan X, each model completed training in approximately 16 hours for 3D models (205 hours for DNA+) and in 7 hours for 2D models. Training of the DNA+ model was extended to 616,880 mini-batch iterations. For prediction tasks, we minimally crop the input image such that its size in any dimension is a multiple of 16, to accommodate the multi-scale aspect of the CNN architecture. Prediction takes approximately 1 second for 3D images and 0.5 seconds for 2D images. Our model training pipeline was implemented in Python using the PyTorch package (http://pytorch.org).

### 3D light microscopy model results analysis and validation

For 3D light microscopy applications, model accuracy was quantified by the Pearson’s correlation coefficient between the model’s output and independent, ground truth test images. To estimate the theoretical upper bound on the performance of a model, we calculated the correlation between a theoretical model which is able to perfectly predict the spatial fluctuations of the signal but is unable to predict the random fluctuations in the target image that arise from fundamentally unpredictable phenomena (such as noise in the electronics of the camera or fluctuations in number of photons collected from a fluorescent molecule). Intuitively as the relative size of random fluctuations increases relative to the size of predictable signal, one would expect the performance of even a perfect model to degrade. The images in Figure 1 of DNA-labeled targets and predictions make this point, in so far as the model can not be expected to predict the background noise in the DNA-labeled imagery. Therefore, to estimate a lower bound on the amplitude of the random fluctuations we analyzed images of cells that were taken with identical imaging conditions but contained no fluorescent labels, for example, images taken with microscope settings designed to detect Hoechst staining, but with cells for which there was no Hoechst dye applied, or images taken with microscope settings designed to detect GFP but with cells with no GFP present. We used the variance of pixel intensities across the image as an estimate of the variance of random fluctuations (*N)*, and then averaged that variance across control images in order to arrive at our final estimate. Calculating the correlation between a perfect model prediction *S* (equal to the predictable image) and an image *T* which is the combination of the predictable image and the random fluctuations (*T*_*x,y,z*_ = *N*_*x,y,z*_ + *S*_*x,y,z*_), is 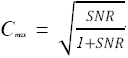 where 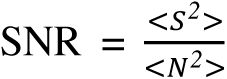. If we assume the correlation between the predictable component and the random component is zero, then the variance of the predictable image (<*S*^2^>) can be calculated by taking the variance of the measured image (<*T*^2^>) and subtracting the variance of the random fluctuations (<*N*^2^>. The result is a formula for the theoretical upper bound of model performance which depends only on the lower bound estimate of the variance of the noise, and the variance of the measured image. We report the average value of *C*_*max*_ for all images in the collection as black tick marks in Figure 1c.

### Registration across imaging modalities

We employed a 2D version of our tool trained on the montage pairs described above. EM images were reflection padded to 1504 px × 1504 px, passed through the trained model, and then predictions were cropped back to the original input size to generate an MBP prediction image. This MBP prediction image was first roughly registered to the larger MBP IF images using cross-correlation-based template matching for a rigid transformation estimate. Next, the residual optical flow^20^ between the predicted image transformed by the rigid estimate and the MBP IF image was calculated, which was then used to fit a similarity transformation that registers the two images, implemented using OpenCV (www.opencv.org). 90 prediction images were randomly selected from the larger set, where more than 1% of the predicted image pixels were greater than 50% of the maximum intensity, to ensure that the images contained sufficient MBP content to drive registration. Ground truth transformation parameters were calculated by two independent authors on this subset of EM images by manual registration (3-4 minutes per pair) to the MBP IF images using TrakEM2. Since the images were registered using a similarity transformation where it is possible for the registration accuracy of the central pixels and those at the edges to be different, the registration errors were calculated by computing the average difference in displacement across an image, as measured in pixels of the target IF image. We report these results for registration differences (Fig. 2) between authors and between the algorithm estimate and one of the authors.

### 3D fluorescence image predictions from a 2D model

To compare performance between models trained on 2D and 3D data, we trained a 2D DNA model for evaluation against the DNA+ model. The 2D model, was trained on the same dataset with the same training parameters as the DNA+ with the exception that training patches of size 64 px × 64 px were sampled from random z-slices of the 3D training images. The model was trained for 250,000 mini-batch iterations with a batch size of 24 for a total training time of approximately 18 hours. After training, 3D predicted fluorescence images were formed by inputing sequential 2D bright-field z-slices into the model and combining the outputs into 3D volumes (Supplementary Fig. 5).

#### Software and Data

Software for training models is available at https://github.com/AllenCellModeling/pytorch_fnet. Data used to train the 3D models is available at https://downloads.allencell.org/publication-data/label-free-prediction/index.html.

## Acknowledgment

We thank the entire Allen Institute for Cell Science team, who generated and characterized the gene-edited hiPS cell lines, developed image-based assays, and recorded the high replicate data sets suitable for modeling and without whom this work would not have been possible. We especially thank the Allen Institute for Cell Science Gene Editing, Assay Development, Microscopy, and Pipeline teams for providing cell lines and images of different transmitted-light imaging modalities, and particularly Kaytlyn Gerbin, Angel Nelson, and Haseeb Malik for performing the cardiomyocyte differentiation and culture, Winnie Leung, Joyce Tang, Melissa Hendershott and Nathalie Gaudreault for gathering the additional time series, CAAX-labeled, cardiomyocyte, HEK-293, and HT-1080 data. We would like to thank the Allen Institute for Cell Science Animated Cell team and Thao Do specifically for providing her expertise in figure preparation. We thank Daniel Fernandes for developing an early proof of concept 2D version of the model. We would like to thank members of the Allen Institute for Brain Science Synapse Biology department for preparing samples and providing images that were the basis for training the conjugate array tomography data. These contributions were absolutely critical for model development. Cardiomyocyte and hiPSC data in this publication were derived from cells in the Allen Cell Collection, a collection of fluorescently labeled hiPSCs derived from the parental WTC line provided by Bruce R. Conklin, at The Gladstone Institutes. We thank Google Accelerated Science for telling us about their studies of 2D deep learning in neurons before beginning this project. This work was supported by grants from NIH/NINDS (R01NS092474) and NIH/NIMH (R01MH104227). We thank Paul G. Allen, founder of the Allen Institute for Cell Science, for his vision, encouragement and support.

## Author Contributions

GRJ conceived the project. CO implemented the model for 2D and 3D images. MM provided guidance and support. CO, SS, FC, and GRJ designed computational experiments. CO, SS, MM, FC, and GRJ wrote the paper.

**Supplementary Figure 1:**
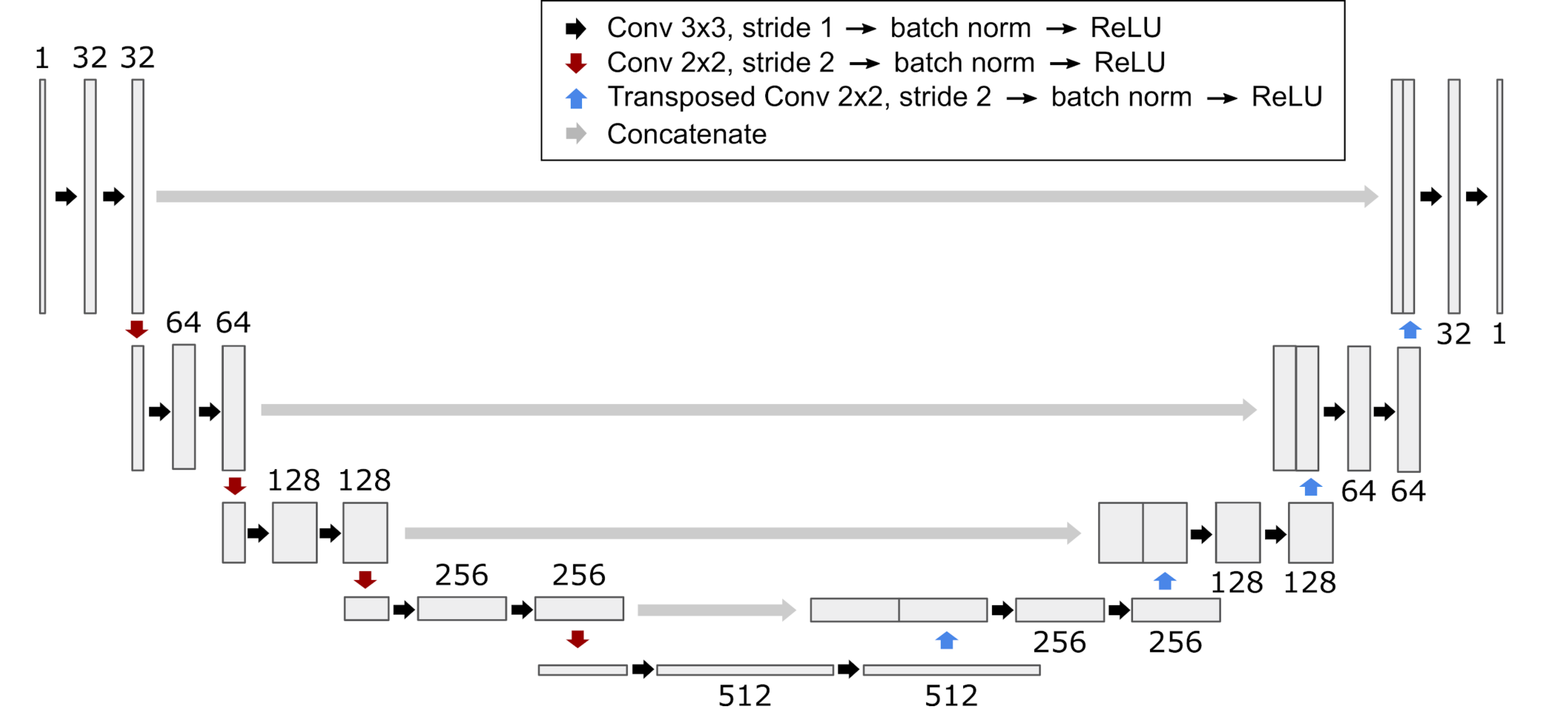
Diagram of CNN architecture underpinning presented tool. There are no batch normalization or ReLU layers on the last layer of the network, and the number of output channels per layer is shown above the box of each layer. Figured adapted from Ronneberger et al.

**Supplementary Figure 2:**
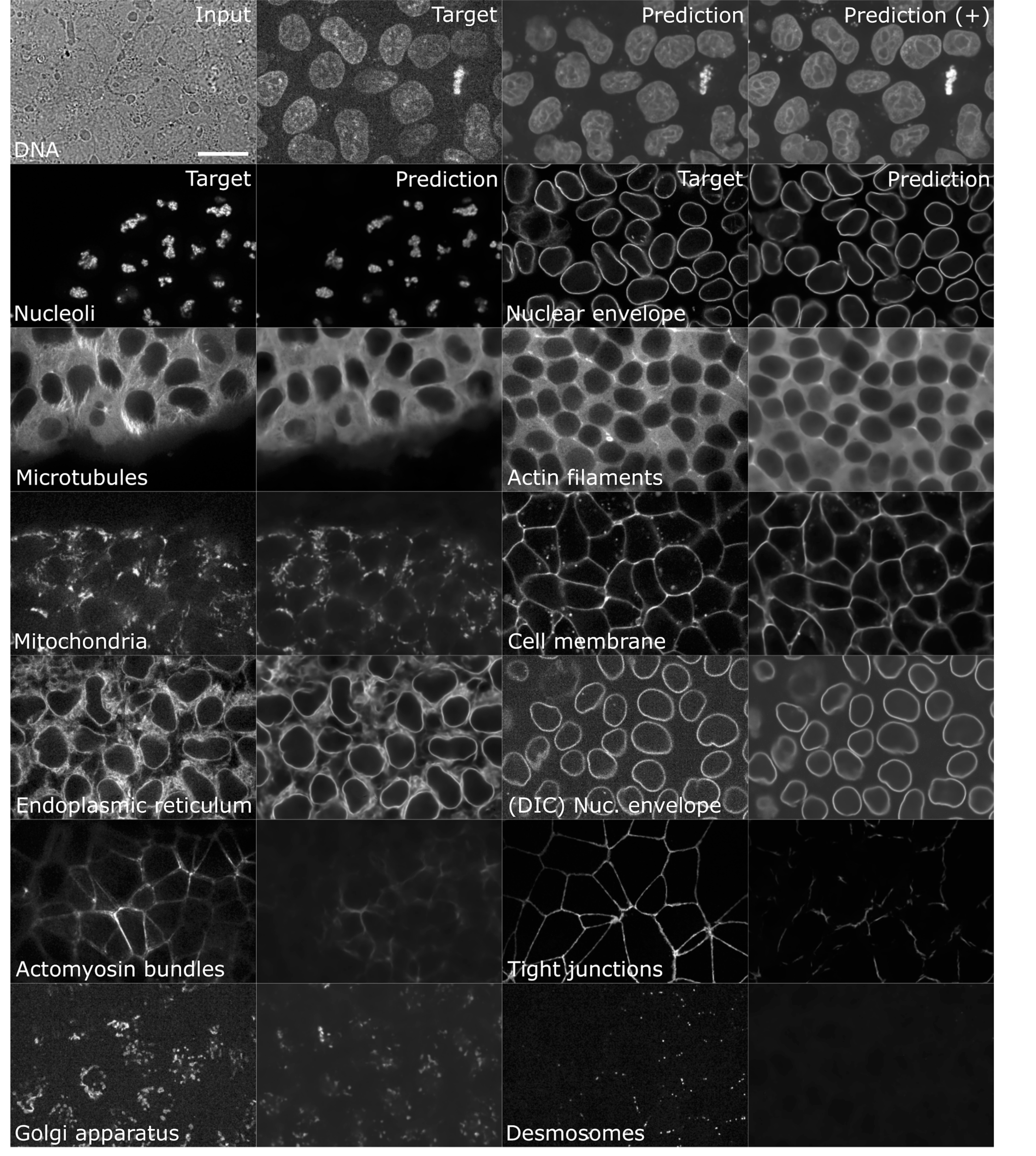
Representative labeled structure models and model predictions for 3D transmitted light microscopy. Results from predicting DNA fluorescence images from DNA and DNA+ models and shown in top row (see Methods). From left, a single z-slice of a 3D transmitted light input image; a ground-truth (“target”, observed) fluorescence image, an image predicted by the DNA model under standard training, and an image predicted by an extended version of the DNA model (DNA+). Subsequent rows are divided into two columns, each with paired images of ground truth and predicted structure localization. In each column, leftmost images show a single z-slice of a ground-truth (“target”, observed) fluorescence image for the labeled structure, while images on the right shown predicted structure localization given standard model training (see Methods)). Left to right and top down presentation order is determined by performance (see Methods, Figure 1c) for nucleoli, nuclear envelope, microtubules, actin filaments, mitochondria, cell membrane, endoplasmic reticulum, nuclear envelope (DIC), actomyosin bundles, tight junctions, Golgi apparatus, and desmosomes models. All models trained on and used bright-field images as inputs (not shown), except where noted (nuclear envelope, DIC). Z-slices were selected to highlight the structure of interest associated with each model. Image-slice pairs were identically contrast stretched, such that black and white values corresponded to the 0.1 and 99.9th percentiles of the target image intensity, respectively. All images shown are independent from model training data. Scale bar is 20 *μ*m.

**Supplementary Figure 3:**
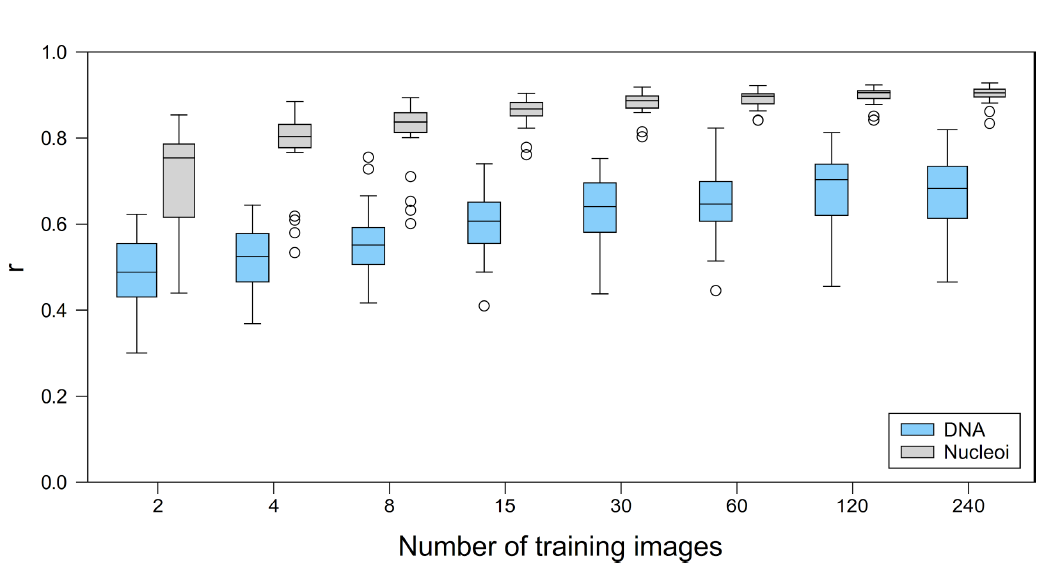
Performance of 3D fluorescence image prediction from transmitted light microscopy (bright-field) image inputs, assessed as a function of number of training images for DNA and nucleoli models. The image-wise correlation coefficient (*r*) was measured between 20 ground-truth (“target”) and predicted fluorescence images. Each target/predicted image pair in the test set is a point in the resultant *r* distribution; the 25th, 50th and 75th percentile image pairs are spanned by the box for each indicated structure, with whiskers indicating the last data points within the 1.5x interquartile range of the lower and upper quartiles. Outliers for each distribution are shown as circles.

**Supplementary Figure 4:**
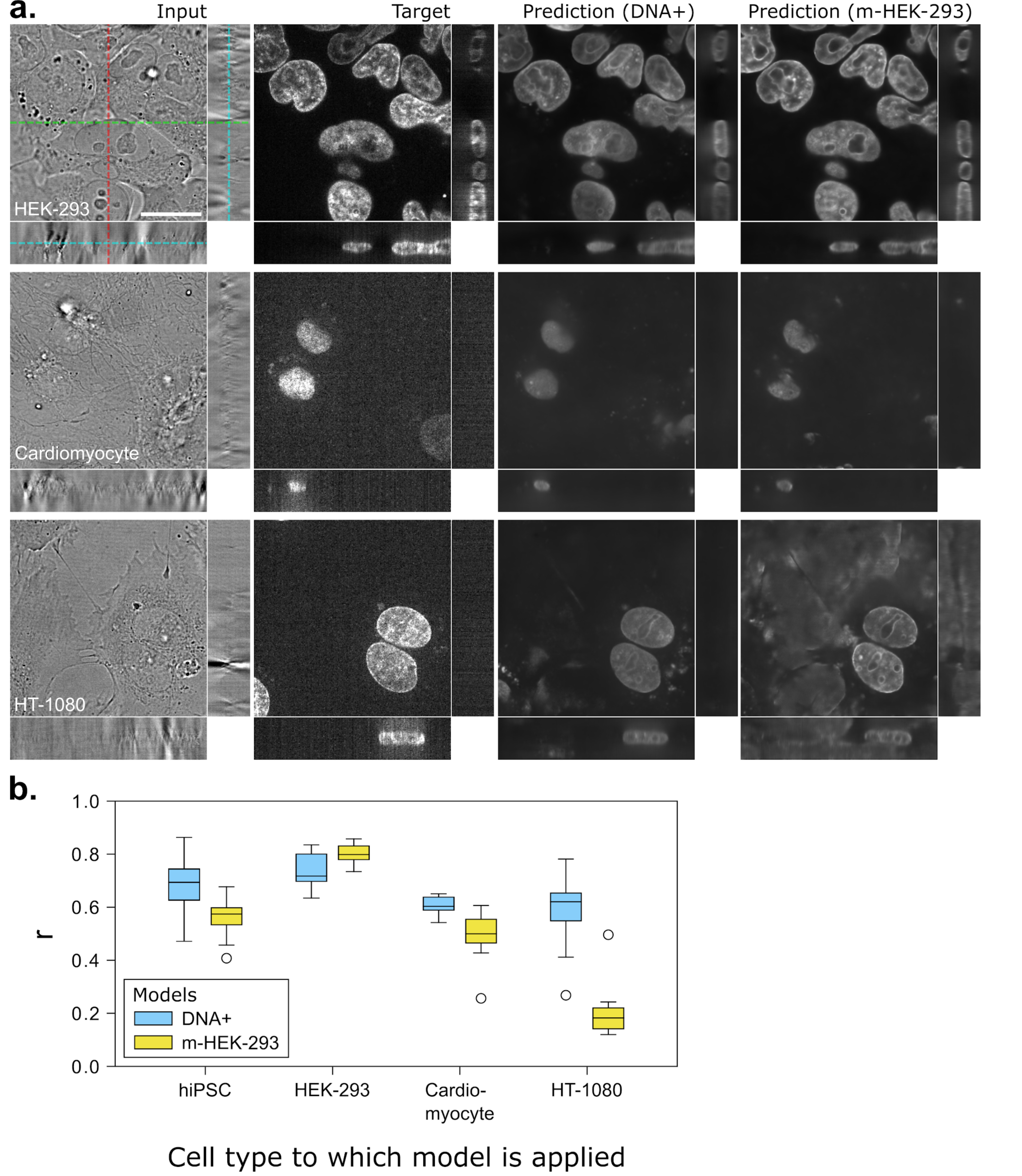
Performance of 3D fluorescence image prediction from transmitted light microscopy (bright-field) image inputs of different cell types. a) Representative DNA predictions on 3D bright-field input images of, from top, HEK-293 cells, cardiomyocytes, and HT-1080 cells using a DNA model trained on hiPSC images (model predicting DNA with an extended training procedure, “DNA+”, see Methods) and a DNA model trained on HEK-293 images (“m-HEK-293”). From left to right, columns are the bright-field input image, the target fluorescence image, the predicted fluorescence image using the DNA+ model, and the predicted fluorescence images using the m-HEK-293 model. The center z-, y-, and x-slices are shown for each 3D image (denoted by the cyan, green, and red lines, respectively, in the left-most image). Scale bar is 20 *μ*m. b) Distributions of the image-wise correlation coefficient (*r*) between target and predicted test images of different cell types using DNA+ and m-HEK-293 models. Each target/predicted image pair in the test set is a point in the resultant *r* distribution; the 25th, 50th and 75th percentile image pairs are spanned by the box for each indicated structure, with whiskers indicating the last data points within the 1.5x interquartile range of the lower and upper quartiles. Outliers for each distribution are shown as circles. Box colors for each input images cell type indicate which model was applied, DNA+ or m-HEK-293. For more details see Methods.

**Supplementary Figure 5:**
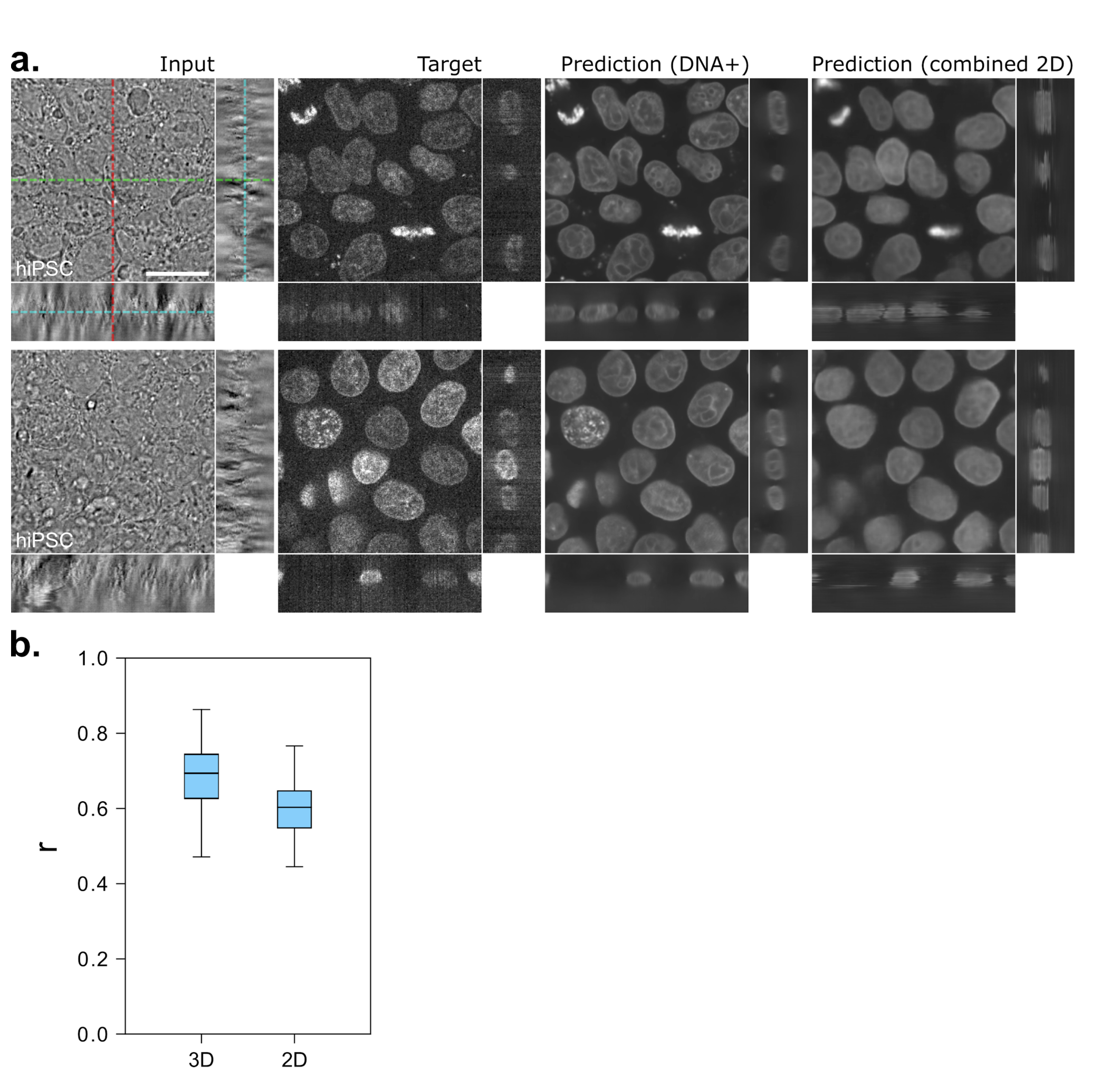
Performance of 3D DNA prediction from transmitted-light (bright-field) images using a fully 3D vs. 2D model. a) Top and bottom rows each show an example of, from left to right, an input bright-field image, a ground-truth (“target”) fluorescence image, a fluorescence image predicted using a 3D model (“DNA+”, see Methods), and a fluorescence image predicted using a 2D model (“combined 2D”, see Methods). 2D model output is composed of predictions on individual input bright-field z-slices combined into a 3D volume. The center z-, y-, and x-slices are shown for each 3D image (denoted by the cyan, green, and red lines respectively in the top-left-most image). Scale bar is 20 *μ*m. The 3D model produces outputs with more detail and accuracy. b) Distributions of the image-wise correlation coefficient (*r*) between target and predicted test images from the 3D and 2D models across 20 test images. Each target/predicted image pair in the test set is a point in the resultant *r* distribution; the 25th, 50th and 75th percentile image pairs are spanned by the box for each indicated structure, with whiskers indicating the last data points within the 1.5x interquartile range of the lower and upper quartiles. Outliers for each distribution are shown as circles.

**Supplementary Figure 6:**
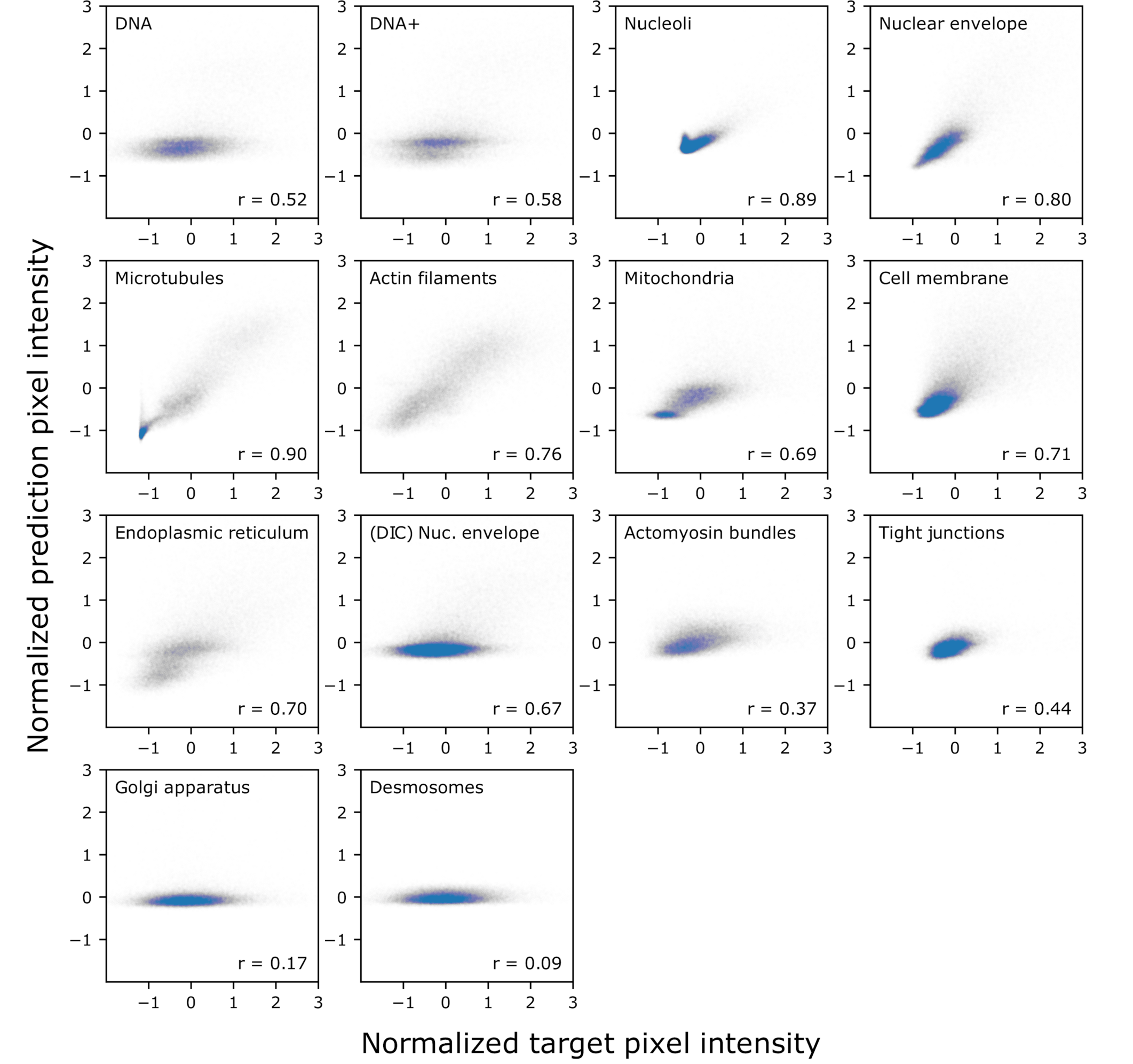
Comparison of ground-truth (“target”) and predicted pixel values in images from Supplementary Figure 2. For each structure indicated, 1% of pixels were randomly selected from the target image are plotted against the corresponding pixels in the predicted image. Pixel intensities are normalized in the target in predicted images (see Methods).

**Supplementary Video 1:**
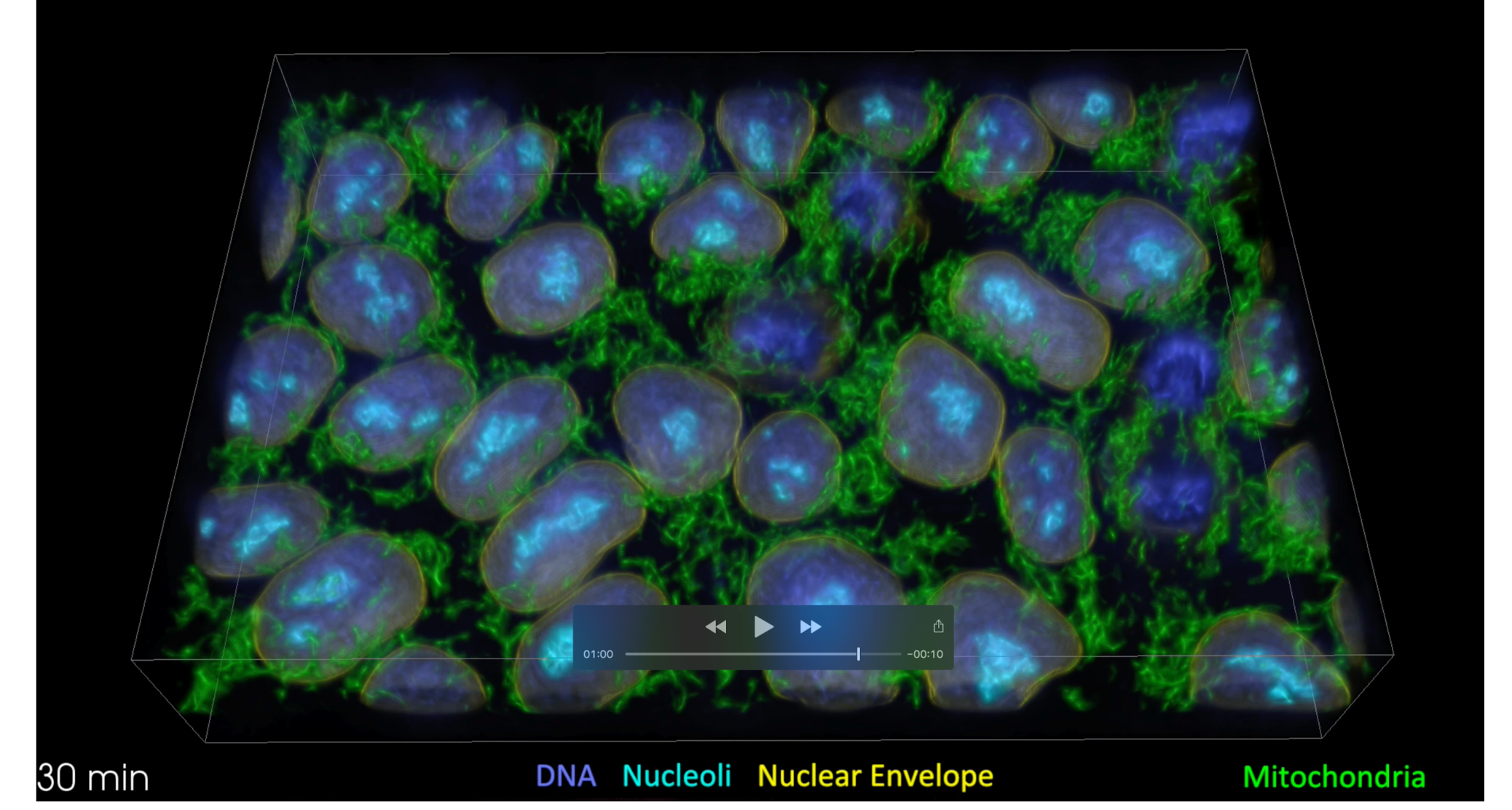
3D rendering of transmitted light microscopy prediction results. Movie illustrates the relationship between 3D time lapse transmitted light (bright-field) input and multiple prediction images. First, individual z-plane images from a 3D transmitted light are shown in succession. Next, individual predictions are shown overlaid in color in the following order: DNA (blue), nucleoli (cyan), nuclear envelope (yellow), cell membrane (magenta), and mitochondria (green). Next, a composite rendering of all channels is shown, followed by a time lapse of single plane from the dataset (also shown in part in Fig. 1e). Finally, a volumetric 3D rendering is shown and played through the individual timepoints 4 times, alternating between showing mitochondria and membrane, together with the nuclear structures (DNA, nuclear membrane, and nucleolus). The boxed outline depicts the extent of the field of view of this volume, which encompasses 97 *μ*m × 65 *μ*m × 19 *μ*m.

**Supplementary Table 1.**
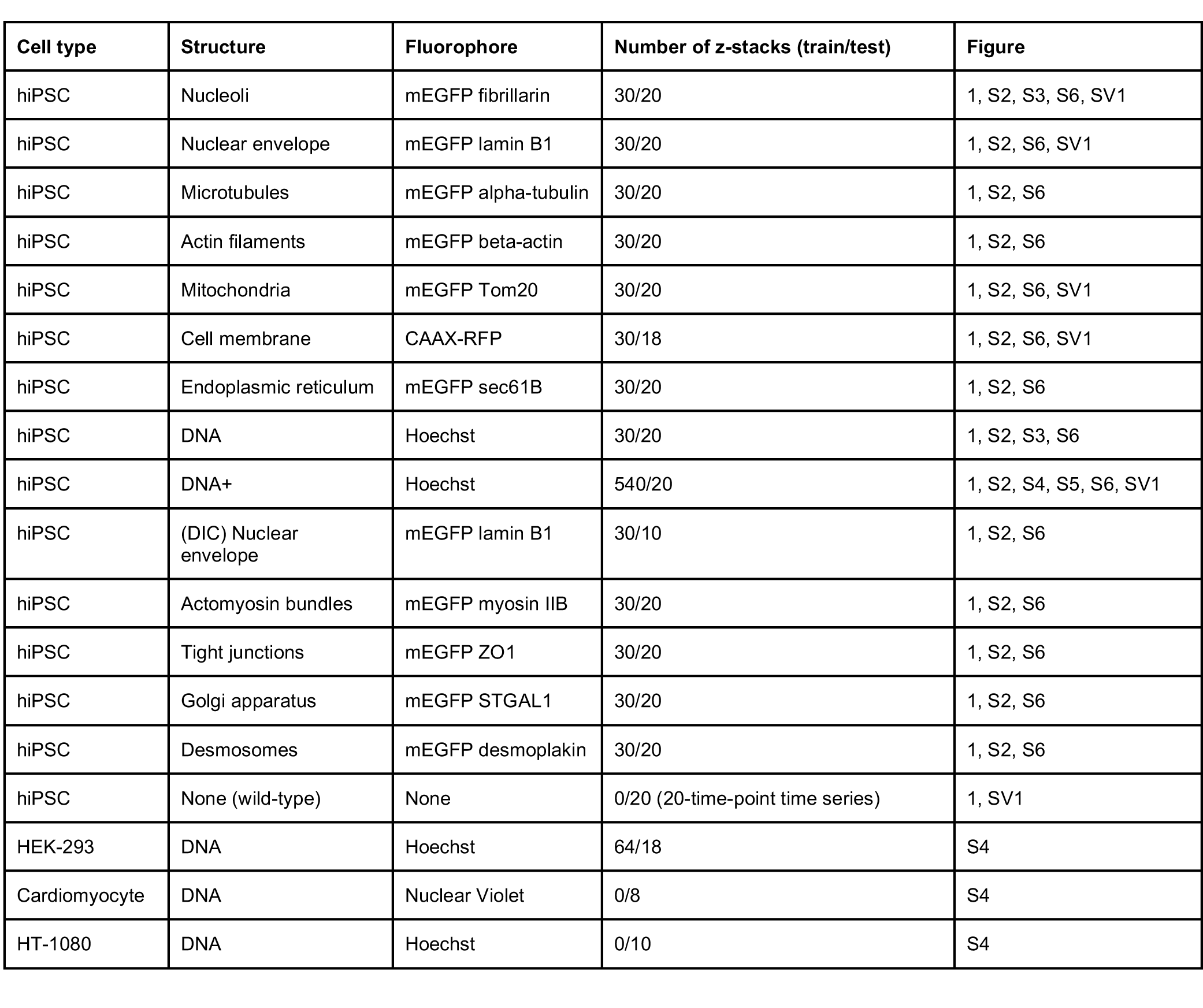
3D live cell imaging data used in this study.

